# Gene Regulatory Divergence Between Locally Adapted Ecotypes in Their Native Habitats

**DOI:** 10.1101/145805

**Authors:** Billie A. Gould, Yani Chen, David B. Lowry

## Abstract

Local adaptation is a key driver of ecological specialization and the formation of new species. Despite its importance, the evolution of gene regulatory divergence among locally-adapted populations is poorly understood, especially how that divergence manifests in nature. Here, we evaluate gene expression divergence and allele-specific gene expression responses for locally-adapted coastal perennial and inland annual accessions of the yellow monkeyflower, *Mimulus guttatus*, in a field reciprocal transplant experiment. Overall, 6765 (73%) of surveyed genes were differentially expressed between coastal and inland habitats, while 7213 (77%) were differentially expressed between the coastal perennial and inland annual accessions. Further, 18% of transcripts had significant genotype x site (GxE) effects. Habitat-specific differential expression was found for 62% of the GxE transcripts (differential expression in one habitat, but not the other), while only 94 (∼5%) GxE transcripts had crossing reaction norms. *Cis*-regulatory variation was pervasive, affecting 79% (5532) of differentially expressed genes. We detected *trans* effects for 52% (3611) of differentially expressed genes. Consistent with the supergene hypothesis of chromosome inversion evolution, a locally adaptive inversion was enriched for *cis*-regulatory divergence. These results provide multiple new insights into the evolution of transcriptome-wide gene regulatory divergence and plasticity among locally adapted populations.

## INTRODUCTION

One of the most important forces driving the evolution of ecological specialization and the origin of biodiversity is local adaptation (Clausen 1951; Colosimo et al. 2005; Sobel et al. 2010). Local adaptation is characterized by reciprocal home-site advantage, whereby populations perform best in their home habitat while performing poorly in foreign habitats (Kawecki & Ebert 2004; Hoekstra et al. 2006; Hereford 2009; Wadgymar et al. 2017). Numerous reciprocal transplant experiments have identified locally adapted populations (Leimu & Fisher 2008; Hereford 2009). Recently, reciprocal transplant experiments have been combined with genetic studies to quantify the role of individual loci adaptive divergence between local populations (Verhoeven et al. 2008; Gardner & Latta 2006; Lowry et al. 2009; Lowry & Willis 2010; Anderson et al. 2011, 2013; Leinonen et al. 2013; Ågren et al. 2013; Wadgymar et al. 2017). Despite this advance of integrating transplant experiments with modern genetic techniques, we have a very limited understanding of how gene expression divergence manifests between locally adapted populations in nature.

One very promising route for understanding how gene expression regulation evolves between locally adaptive populations is to conduct allele specific expression (ASE) analyses in natural populations (Tung et al. 2009; Lovell et al. 2016; Wang et al. 2017). By simultaneously evaluating different causes of gene expression, ASE analyses can determine the relative contributions of variation in nearby *cis* elements versus distant *trans* factors on gene expression divergence (Cowles et al. 2002; Wittkopp et al. 2004). Conducting ASE analyses in conjunction with reciprocal transplant experiments offers the opportunity go beyond partitioning gene expression into *cis* and *trans* effects by establishing how habitat context influences those sources of variation in gene expression. ASE analysis is often conducted in F1 hybrids, where parental alleles are evaluated for significant differences in expression (de Meaux et al. 2005; Guo et al. 2008; Zhang & Borevitz 2009; Cubillos et al. 2014; Coolon et al. 2014; Steige et al. 2015, 2017; Lovell et al. 2016; Aguilar-Rangel et al. 2017). ASE relies on there being nucleotide polymorphisms between parental alleles, so that those alleles can be individually counted in the F1 transcript pool. *Cis*-regulatory allelic expression differences are expected to persist regardless of whether alleles are expressed separately in parental genotypes or together in an F1 hybrid. In this way, ASE allows for the identification of *cis* expression Quantitative Trait Loci (eQTL), which can be evaluated for their responses to different environmental conditions (Lovell et al. 2016). Further, *trans* effects on gene expression are indicated by a change in allelic effects between the parental generation and the F1 hybrids (Cowles et al. 2002; Wittkopp et al. 2004). Despite its great potential, we are unaware of any field reciprocal transplant experiment that has measured ASE to evaluate how gene expression divergence manifests across the habitats known to be responsible for local adaptation.

In this study, we evaluated gene regulatory divergence in nature between locally adapted coastal perennial and inland annual population accessions of the yellow monkeyflower, *Mimulus guttatus* (*syn. Erythranthe guttata* (Fisch. ex DC.) G.L. Nesom). *M. guttatus* is a model system for understanding the genetic underpinnings of evolutionary adaptations (reviewed in Wu et al. 2008; Twyford et al. 2015). Both annual and perennial populations are found throughout much of the range of *M. guttatus* (van Kleunen 2007; Lowry & Willis 2010; Friedman & Willis 2013; Oneal et al. 2014; Twyford et al. 2015). The large differences in morphology and life-history among populations of *M. guttatus* has been shown to be genetically-based through common garden experiments (Vickery 1952; Hall & Willis 2006; Lowry et al. 2008; Lowry & Willis 2010). In the inland Mediterranean climates of California’s Coast Range and Sierra Foothills, most populations of *M. guttatus* have an early flowering annual life-history, which they have evolved to escape from the hot summer drought endemic to these regions. In contrast, coastal populations of *M. guttatus* are sheltered from the summer drought by pervasive marine fog, which supplies soil moisture and reduces transpiration rates (Corbin et al. 2005). As a result, coastal populations uniformly have a late-flowering perennial life-history (Hall & Willis 2006; Lowry et al. 2008). While coastal habitats are benign in terms of soil water availability, these populations must contend with toxic oceanic salt spray, in response to which they have evolved high salt tolerance (Lowry et al. 2008, 2009; Selby et al. 2014). Across previous reciprocal transplant field experiments, inland plants survived to flower at 23.9 times the rate of coastal plants in inland habitat, while coastal plants have survived at 2.5 times the rate of inland plants in coastal habitat (Hall & Willis 2006; Lowry et al. 2008, 2009; Hall et al. 2010).

Population structure analysis has repeatedly demonstrated that coastal populations, all of which are perennial, collectively constitute a single ecotype that is genetically distinct from inland populations of the species (Lowry et al. 2008; Twyford & Friedman 2015). The individual genes responsible for the divergence of coastal perennial and inland annual populations are unknown, but it is clear that a large paracentric chromosomal inversion on chromosome 8 is responsible for much of the divergence in traits and fitness between these populations (Lowry & Willis 2010; Friedman 2014; Oneal et al. 2014; Gould et al. 2017). Smaller effect loci across the genome also account for divergence between the coastal perennial and inland annual populations (Hall et al. 2006, 2010; Lowry et al. 2009; Friedman et al. 2015). The chromosome 8 inversion is also involved in the divergence of inland annual populations from inland perennial populations (Lowry & Willis 2010). Divergence of inland annual and perennial populations was not a focus of this study.

The primary goal of our study was to characterize gene regulatory divergence between coastal perennial and inland annual *M. guttatus* in the natural habitats that drove the evolution of local adaptation among these ecotypes. In particular, our study aimed to 1) determine the prevalence of genotype x environment (GxE) interactions affecting the expression of native and non-native alleles across the divergent habitats, 2) characterize the relative contributions of *cis* and/or *trans* regulatory variation in expression divergence, 3) establish the role of genome structure (e.g. inversions) in the evolution of potentially locally adaptive regulatory variation, and 4) identify candidate genes underlying local adaptation in this system. The results of our study provide new insights into the gene regulatory evolution and suggest that coupling allele-specific expression with reciprocal transplant experiments is a promising approach for modern local adaptation research.

## MATERIALS AND METHODS

### Germplasm

Original field collections of the seeds used in this experiment were made in the Spring of 2005 from the SWB (N39.0359’, W123.6904’) and LMC (N38.8640’, W123.0839’) populations located in Mendocino County, California. The SWB and LMC populations have been the subject of extensive research since then (Lowry et al. 2008; Lowry & Willis 2010; Wu et al. 2010; Friedman & Willis 2013; Friedman 2014; Gould et al. 2017). Seeds from these populations were inbred for multiple generations in the Duke University Greenhouses. The plants used for the field experiment included the following: 1) An inbred (7 generations self-fertilized) inland line, LMC-L1-2, henceforth referred to as LL; 2) An inbred (13 generation self-fertilized) coastal line, SWB-S1-2, henceforth referred to as SS; and 3) An F1 hybrid between the two inbred lines, henceforth referred to as F1 or SL. The F1s used in this experiment were made in the Michigan State University (MSU) Growth Chamber Facilities through a cross between the two parental inbred lines.

### Field Reciprocal Transplant Experiment

We conducted a field reciprocal transplant experiment in tandem with allele-specific expression analyses to understand the causes (*cis, trans*) of gene expression divergence between locally adapted coastal perennial and inland annual populations of *M. guttatus* (Fig. 1). The reciprocal transplant experiment was conducted in 2016 using one coastal (Bodega Marine Reserve; N38.3157’, W123.0687’) and one inland (Pepperwood Preserve; N38.5755’, W122.7008’) field site located in Sonoma County, CA. SS, LL, and F1 seeds were originally sown in Sun Gro Metromix 838 soil on February 3. The seeds were then stratified in the dark at 4°C until February 9, when they were removed from the cold and germinated in the UC Berkeley Oxford Track Greenhouses. Seedlings were moved to the Bodega Marine Reserve Greenhouses on February 21 for acclimation. On February 22, the seedlings were transferred to native soil collected from the field sites. Seedlings were planted in the field at Bodega on March 7 and at Pepperwood on March 8. ∼90 plants per type (SS, LL, F1) were planted at each field site. Plants were evenly distributed among three blocks per field site and randomized within block. Tissue for RNA sequencing was collected from Bodega on April 23 and Pepperwood on April 24. To avoid effects of circadian rhythm on gene expression, all tissue was collected between 12:00pm and 1:00pm. Both collection days had mostly clear skies, with mean local temperatures during collection periods of 14° C at Bodega and 17° C at Pepperwood. Plants had progressed to different developmental stages at the time of collection, with some plants approaching flowering and others not. To best standardize tissue collection across genotypes, only the two youngest pairs of fully expanded leaves were collected from each plant. Tissue was immediately flash frozen on liquid nitrogen in the field. Samples were then shipped on dry ice to Michigan State University for RNA extractions.

**Fig. 1.**
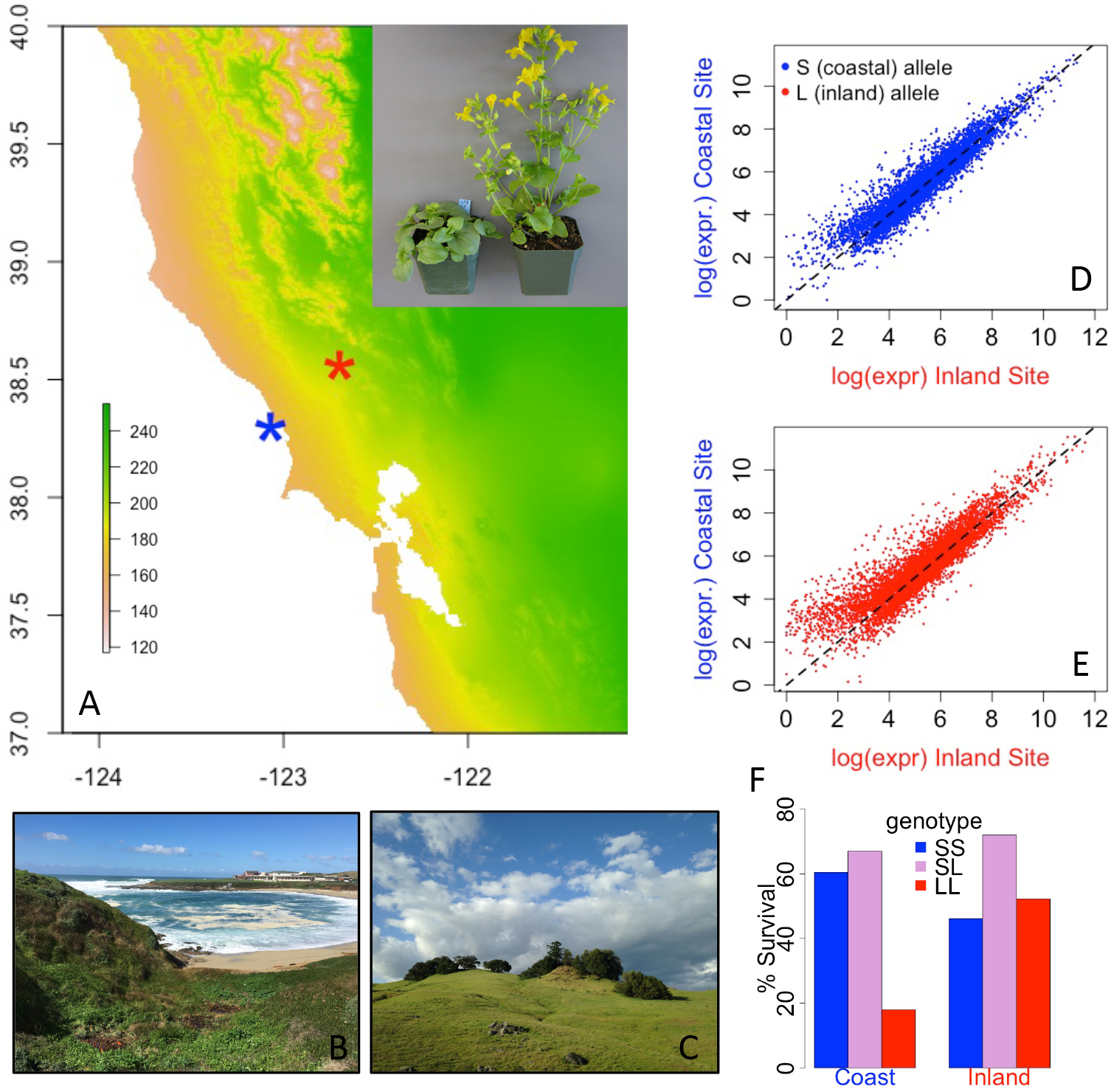
A) Map of Northern California with the location of two field transplant sites. Colors represent mean temperature of the warmest quarter in degrees C x 10. Inset: left, coastal ecotype; right, inland ecotype. B) The coastal field site. C) The inland field site. D) SS parental gene expression at two field sites. E) LL parental gene expression at two field sites. F) Survival of parental and F1 (SL) plants at the two field sites.

### RNA-sequencing

Total RNA was extracted from 80 leaf tissue samples with the Spectrum Plant Total RNA Kit (Sigma-Aldrich, St. Louis, MO, USA). RNA was extracted from 12 replicates of both SS and LL parental lines from the coastal field site and 13 replicates of both parental lines from the inland field site. RNA from 15 F1 hybrids from each site was also extracted. Total RNA was submitted for library preparation and sequencing at the MSU Genome Core (https://rtsf.natsci.msu.edu/genomics/). RNA-Seq libraries were prepared using the Illumina TruSeq Stranded mRNA Library Preparation Kit with a Perkin Elmer (Waltham, MA) Sciclone NGS robot. Completed libraries were quality controlled and quantified using a combination of Qubit dsDNA HS (Thermo Fisher Scientific, Waltham, MA), Caliper LabChipGX HS DNA (Perkin Elmer, Waltham, MA), and Kapa Biosystems Illumina Library Quantification qPCR (Wilmington, MA) assay kits. All libraries were barcoded and library concentrations were normalized and pooled in equimolar amounts: 8 pools each consisting of 10 randomly chosen libraries were made. Each of these pools was loaded on one lane of an Illumina HiSeq 2500 High Output flow cell (v4) and sequenced in the 1×100bp (SE100) format using HiSeq SBS reagents (also v4). Base calling was done by Illumina Real Time Analysis (RTA) v1.18.64 and output of RTA was demultiplexed and converted to Fastq format with Illumina Bcl2fastq v1.8.4.

### Reference Transcriptome Assembly

Alignment of RNA-seq to a single reference genome can lead to major biases in ASE studies (Stevenson et al. 2013). To overcome reference bias, we assembled reference transcriptomes for the coastal perennial (SS) and inland annual (LL) parents. We chose this method over construction of a pseudo-transcriptome reference (Shen et al. 2013) to avoid generating bias caused by the reference genome and transcriptome for *M. guttatus*, which was assembled from an inland annual line (IM62; Hellsten et al. 2013). Reference bias is of particular concern in *M. guttatus* because pairwise polymorphism is generally high (π_S_ = 0.033-0.065; Brandvain et al. 2014; Gould et al. 2017; Puzey et al. 2017). To generate a reference transcriptome assembly that contained allele-specific sequences for each parental genotype, we sequenced and conducted *de novo* assembly using RNA-sequencing. Plants of each of the coastal (SS) and inland (LL) lines were grown in the MSU greenhouses until both genotypes were flowering. Floral bud, leaf, and root tissue were collected from each of the SS and LL lines, flash frozen with liquid nitrogen, and stored at −80° C. Total RNA was extracted from each sample using the Spectrum Plant Total RNA Kit and submitted to the MSU Genome Core for library construction and sequencing. Strand-specific Tru-Seq Illumina RNA-seq libraries were prepared and sequenced along with four libraries constructed from field collected leaf tissue from the reciprocal transplant experiment (one sample per parental line per field site). These libraries were included in the transcriptome assembly pipeline to capture sequences from genes that may only be expressed under field conditions. Completed libraries were quality controlled and quantified using a combination of Qubit dsDNA HS, Caliper LabChipGX HS DNA, and Kapa Biosystems Illumina Library Quantification qPCR assay kits. A total of 10 libraries were barcoded with dual index barcodes and pooled together. The entire pool was sequenced using 125 bp paired-end reads on a single lane of the Illumina GA 2500 platform.

Raw output reads were quality checked using FastQC v.0.11.3 (Andrews 2015). Residual adapter sequences were removed and reads were quality trimmed using Trimmomatic v.0.33 (Bolger et al. 2014), discarding sequences less than 50 bp in length. Paired and un-paired reads from each ecotype were combined and assembled into two separate *de novo* (non-reference guided) assemblies using Trinity v20140413p1 (Grabherr et al. 2011) with *in silico* read normalization. To check the quality of our two transcriptomes, we compared our *de novo* assemblies to the set of primary transcripts from the IM62 reference transcriptome using Trinity utility scripts. We measured how much of each *Mimulus* primary transcript was reconstructed in the two assemblies. We found that 74.3% of *Mimulus* primary transcripts were reconstructed (to at least 50% of their full length) in the inland parent (L) transcriptome. In the coastal parent transcriptome, 74.1% of *Mimulus* primary transcripts were reconstructed (to at least 50% of their full length). Thus, the two assembled transcriptomes were of similar quality and completeness. We aligned the filtered input reads back to the finished assemblies using Bowtie2 (Langmead et al. 2012). A total of 98.98% and 99.26% of input reads aligned back to the inland (L) and coastal (S) assemblies, respectively. We identified possible non-plant contaminant sequences in each assembly through comparison to the NCBI nt database using blastn. We eliminated transcriptome sequences with top blast sequence matches (e-value cutoff of 1e-20) to non-plant organisms (*N* = 16,243 and *N* = 1,222 eliminated sequences for the inland and coastal transcriptomes, respectively).

To construct a combined reference transcriptome that contained pairs of allelic sequences from both lines, we used a reciprocal best blast strategy. Each transcriptome was compared against the other (blastn) at an e-value cutoff of 1e-5 and reciprocal best hits were returned using a custom Python (v. 2.7.2) script (https://github.com/lowrylab/Mimulus_Gene_Expression). We tested a range of e-value cutoffs in this analysis and found that the choice of e-value between 1e-5 and 1e-50 had little effect on the total output number of best blast pairs (see Fig. S1). We considered two sequences best blast hits if total high-scoring pair (HSP) coverage was 75% or greater between sequences. We included only aligned sequence regions of the longest HSP in the combined transcriptome.

To match transcripts in the combined *de novo* reference transcriptome with their corresponding *Mimulus* genes, we compared *de novo* assembled alleles against the reference (IM62) set of primary transcripts (blastn) at an e-value cut-off of 1e-5. In rare cases, where two alleles from a *de novo* assembled transcript matched different *Mimulus* genes, the gene with the lowest e-value alignment was chosen as a match for the allele pair. Note that with this annotation method, each *de novo* assembled transcript has only one matching gene and thus, can be unambiguously assigned a specific location in the genome. However, multiple transcripts can be assigned to the same genomic location, which is typical of isoforms stemming from the same gene.

### Differential Expression Analyses

Raw reads from the 80 field RNA-seq libraries were de-multiplexed, quality filtered, and trimmed using FastQC and Trimmomatic, as described above for the transcriptome assembly. Filtered reads were aligned to the *de novo* combined reference transcriptome (described above) using the un-gapped aligner Bowtie v1.0.0 (Langmead et al. 2009). We outputted only unique alignments with zero mismatches. In this way, we analyzed only read count data for reads that aligned uniquely and perfectly to only one parental allele in the combined reference transcriptome. Only reads that overlap one or more polymorphisms between the parental alleles provide useful information on differential expression between alleles. Approximately 52% of reads per individual had one or more than one alignment to the transcriptome. Unaligned reads fall mainly in areas that were not reconstructed in the reference transcriptomes. On average 28.6% of library reads per individual aligned uniquely to only one allele and were thus, informative for allele-specific expression (range 23.7 – 30.7%). For confirmation, some alignments were visually checked using IGV v.2.3.65 (Thorvaldsdóttir et al. 2013).

Read alignments to each transcript were quantified using a custom Python (v.2.7.2) script. Using the raw count data, we identified allele pairs in the transcriptome for which there was a high probability of residual heterozygosity in one of the parental inbred lines. Potential regions of residual heterozygosity were indicated when a significant number of reads from a homozygous parent aligned to the opposite parental allele in the transcriptome. We eliminated from further analysis transcripts where the average number of reads aligning to the correct parental allele was less than 5 times the average number of reads aligning to the alternate allele (*N* = 2255 eliminated transcripts). For example, we eliminated transcripts where the SS parental line did not have at least 5 times more reads aligning to the S allele than the L allele in the reference transcriptome. We also used the initial count data to test for mapping bias (Fig. S2). For both field sites, mapping bias was present but well controlled. On average there were 1.9% more reads aligned to the S allele than the L allele for F1 plants (Fig. S3). There was a small but statistically significant relationship between transcript nucleotide divergence and differential expression (equivalent to 0.001 increase in the absolute value of LFC between genotypes, per SNP). Count data was normalized for library size using Trimmed Mean of M-values (TMM) normalization to counts per million, which reduces biases generated by the presence of very highly and lowly expressed transcripts (Robinson et al. 2010). We eliminated any transcript without at least one library containing >= 10 counts per million reads. The remaining sequences constitute the ‘expressed’ set of transcripts (*N* = 10,122). The mean expressed library size for samples from the coast and inland sites was not significantly different (*t*-tests: all libraries, *P* = 0.60; parental libraries only, *P* = 0.22, Fig. S4, and is thus, unlikely to strongly affect statistical comparisons between the two sites. Only 2 transcripts were expressed at only one site and only 7 transcripts were expressed by only one parental line.

We analyzed the expressed transcript data using the limma (Ritchie et al. 2015) and edgeR (Robinson et al. 2010) packages in R v.3.2.1 (R. Core Team 2012). Variance stabilizing normalization of the data was performed using the voom (Law et al. 2014) function in limma, with a design matrix corresponding to the particular linear model being used for analysis. Model design matrices were coded using the default treatment-contrast parameterization scheme (see Smyth et al. (2015), section 9.5.4) and the significance of model effects were calculated using voom precision weights and the eBayes function. Model effects were judged significant if they fell below a Benjamini-Hoechberg false-discovery rate corrected P-value of 0.05 using the decideTests function.

We used several different linear models to analyze the count data for expressed transcripts, modified from the method of Lovell et al. (2016). For this approach, it is important that the size of the L and S sub-transcriptome in each F1 is approximately equal to produce unbiased results, which we verified independently (Fig. S3). Each linear model had one or more fixed factors: allele (L/S), environment (coastal field site vs. inland field site), generation (F0/F1), and interaction terms. We analyzed the parental plants (F0 generation) separately, the F1 plants separately, and both generations together in different models. We avoided analyzing models with more than one categorical factor, except for the purposes of measuring interaction effects because effect significance testing for crossing categorical factors only captures differences present in the first level of each factor.

For data from the F0 plants, we ran models examining the allele (homozygous genotype) effect within each environment separately. We also tested for environmental effects separately within each parental genotype. We used a model with both genotype and environment to test the significance of the genotype x environment interactions. Because the inland site had one extra plant per genotype compared with the coastal site, we analyzed data from 12 randomly chosen parental lines of each genotype to equalize the power to detect effects between sites. A *post-hoc* power analysis was conducted using the R package *RnaSeqSampleSize* (Bi & Liu 2016). We used data from all plants together (both F0 and F1) to test for *cis* and *trans* regulation. *Cis*-regulatory allelic expression differences are expected to persist regardless of whether alleles are expressed separately in parental genotypes or together in an F1 hybrid. However, in the F1 genotype, *trans* acting factors from both parental genotypes will influence the expression of an allele, so *trans* regulation is indicated if allelic patterns of differential expression are different in the parents vs. the F1 plants. In our model, a significant effect of allele is indicative of a *cis*-regulatory difference in expression. Within each environment, we tested a model with the following factors: allele (genotype), generation (F0/F1), and the interaction term. Any transcript with any *cis* regulation had a significant allele effect (measured in the F1 generation only). Any transcript with only *trans* regulation had a significant difference between generations, but a non-significant allele effect in the F1 generation. Any transcript with both *cis* and *trans* regulation had a significant generational effect and a significant allele effect in the F1.

To gauge the significance of *cis* x environment interactions, we tested the interaction term of a model using only F1 data and the factors allele, environment, and interaction. To gauge the significance of trans x environment interactions, we modeled all the data (F0 and F1) and measured the significance of a three-way interaction term: allele x generation x environment.

### Gene Ontology Enrichment and Expression Networks

We tested for enrichment of gene-ontology (GO) terms among subsets of genes with particular expression patterns (significant model effects). To annotate GO terms, all IM62 reference *Mimulus* proteins were paired with their best *Arabidopsis* protein matches (blastp) at an e-value cutoff of 10^-3. The set of expressed transcripts in our study was matched to the IM62 reference genome genes, as described above, and corresponding GO terms were matched to the *Arabidopsis* BLAST hits. Subsequent GO enrichment tests were done using the R BIOCONDUCTOR package topGO (Alexa & Rahnenführer 2015). We used the Bioconductor database ‘org.At.tair.db’ for annotation and the algorithm ‘classic’ (statistic = ‘fisher’) for statistical tests. Because P-value distributions of GO enrichment tests do not typically conform to a distribution amenable to Benjamini-Hoechberg FDR control, and because such control methods can be overly conservative causing loss of valuable functional information (Alexa & Rahnenführer 2015), we report GO terms with uncorrected enrichment *P*-values <= 0.01 (Table S1). Enrichment tests were conducted at the level of genes rather than transcripts. For each gene in the IM62 reference transcriptome, we determined the model effects for all matching expressed transcripts in our data set. A model effect for a gene was considered significant if at least one matching transcript had a significant effect. GO enrichment was then tested among all expressed genes (*N* = 9326) with a significant effect vs. all expressed genes without a significant effect. Enrichments for GxE effects were measured against all genes where either environmental or allelic DE could be detected. Reaction norm plots were generated using the LSMean expression level of all transcripts for each gene in each environment.

We conducted weighted gene co-expression network analysis within each parental genotype using the R package WGCNA (Langfelder et al. 2008). The analysis identifies genes with correlated patterns of expression across individuals (eigengenes), finding gene sets that putatively form molecular networks (or modules). We conducted WGCNA separately for each parental genotype and then identified which networks in each parent had expression that was significantly different between field sites. Gene expression was quantified as the expression level of the longest matching transcript for each gene. For all analyses, we calculated dissimilarity of expression using a soft thresholding power of 6 and used the dynamicTreeCut function to define modules with a minimum number of 30 genes. We calculated the correlation of each cluster (eigengene) with habitat (Fig. S5). Gene ontology term enrichment was calculated for all modules with a correlation value of r > 0.75 (Table S2).

### Chromosomal Inversion Analyses

Two inverted regions of the genome, one on chromosome 5 and one on chromosome 8, have been linked to adaptive differences between inland and coastal ecotypes in previous studies (Lowry & Willis 2010; Oneal et al. 2014; Twyford & Friedman 2015; Gould et al. 2017). When examining genes in putatively inverted regions of the genome, we included reference genome v1.1 scaffolds that are inferred to be in the inverted regions from a previous linkage mapping study (Holeski et al. 2014). For the inversion on chromosome 5, we included v1.1 scaffolds 36, 149, 158, 170, 197, 266, 281, 288, 327, and 368. For the inversion on chromosome 8, we included v1.1 scaffolds 11, 59, 76, 155 233, 604, and 1093. We tested whether genes in these regions of the genome are enriched for any model effects. Enrichment tests of model effects among annotated inversion and non-inversion genes were carried out using Fisher’s Exact tests. The magnitude of model effects among genes with significant effects were compared using Wilcox rank sum tests in R. It should be noted that the presence of alternative orientations of chromosome 8 inversion in the SS and LL parental lines has been confirmed by previous crosses (Lowry & Willis 2010) while the chromosome 5 inversion was originally identified in a cross between different inland and coastal lines (Holeski et al. 2014). High differentiation between the SS and LL parental lines in both inversion regions has been demonstrated by an allele frequency outlier analysis (Gould et al. 2017).

### Patterns of Candidate Gene Expression

In a study of species-wide genomic variation, Gould et al. (2017) identified a set of 667 candidate adaptation genes. These genes had unusually high differentiation in allele frequencies between a set of 47 coastal and 50 inland *Mimulus* populations. High differentiation was detected in either the genic or 1 kb upstream promoter regions of the genes, or both. We tested for enrichment of differential expression patterns among the expressed candidate genes in the current study. Enrichment was tested against all expressed non-candidate genes using Fisher’s Exact tests. The magnitude of model effects among genes with significant effects were compared using Wilcox rank sum tests in R.

## RESULTS

### Reference Transcriptome Assembly

From the tissues of the two *M. guttatus* parental lines we were able to reconstruct both parental alleles for 25,893 transcripts, of which 708 pairs were removed because they contained no polymorphism between lines. Eighty-two percent of transcripts could be matched back to previously annotated *M. guttatus* genes (www.phytozome.org). Our transcriptome reconstructed 54.0% of *M. guttatus* primary transcripts at 50% coverage or greater (Fig S5). On average, there were 2.8 polymorphisms (SNPs or indels) per 100 bp for each pair of alleles in the reference transcriptome (Fig. S6C).

### Differential Expression Analyses

RNA-sequencing of plants from the field experiments produced approximately 25.9 million raw reads per library (range 9.2-35.9 million reads). We were able to evaluate gene expression for 10,122 transcripts corresponding to 9,326 genes, after transcriptome reconstruction and filtering. Significant expression differences (FDR = 0.05) between the inland and coastal field sites were common (6765 genes; 73% of genes). Expression differences between parental genotypes within field sites were also common (7213 genes; 77%; Table 1 and 2). Most transcripts that were differentially expressed between parental lines were also sensitive to environment: only 7.2% of transcripts were consistently differentially expressed between lines regardless of field site. We found that 54% more genes were differentially expressed between parental lines at the inland site than at the coastal site (Table 2). We can attribute only part of this discrepancy to differences in power at the two sites. Average power to detect DE genes was only slightly higher at the inland site than the coastal site (75% vs. 68% for a minimum fold-change of 2X).

**Table 1:**
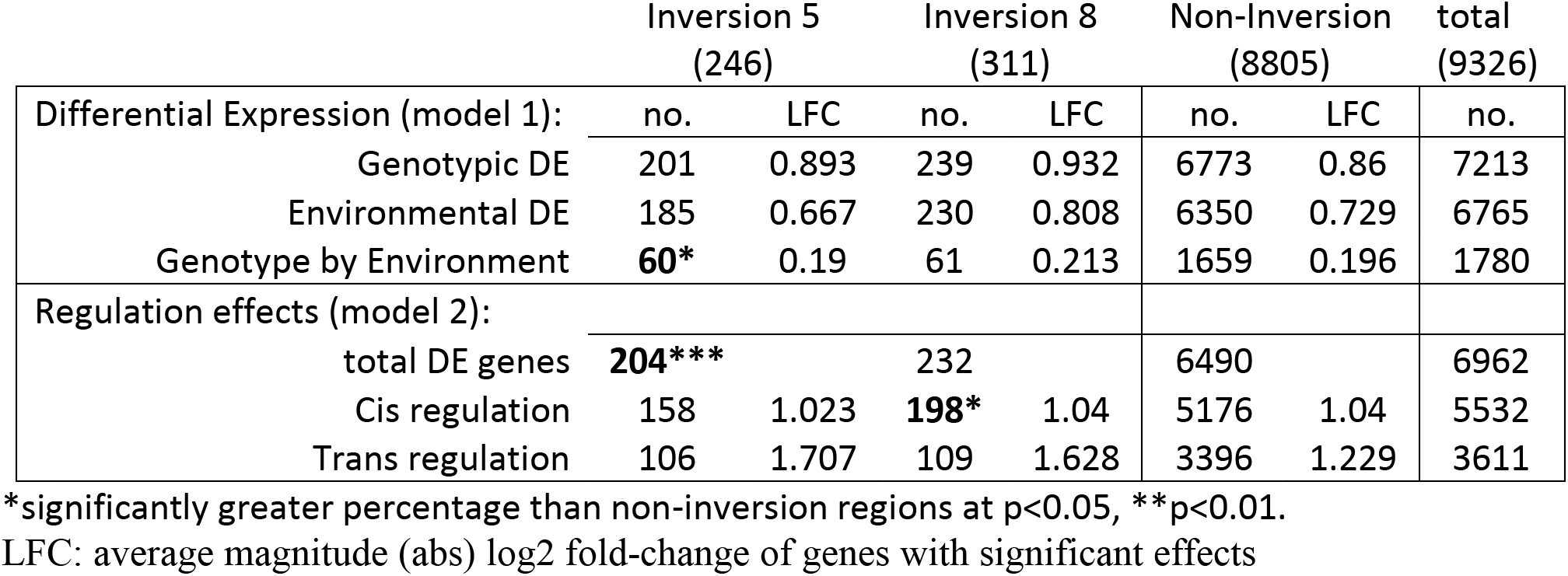
Summary of model effects on expressed genes.

**Table 2:** Number and average effect size of differentially expressed genes within each habitat.

|  | Coast |  | Inland |  |
| --- | --- | --- | --- | --- |
|  | no. | LFC | no. | LFC |
| Inversion 5 (246) | 119 | 1.136 | 184 | 0.932 |
| Inversion 8 (311) | **161\*** | 1.092 | 210 | 0.993 |
| Non-Inversion (8805) | 3997 | 1.084 | 6195 | 0.897 |
| Total (9326) | 4277 | 1.085 | **6589\*\*\*** | **0.901\*\*\*** |

\*significantly greater than non-inversion regions at $p < 0.05$
\*\*\* significantly different than the coastal field site, $p < 0.001$
LFC: average magnitude (abs) log2 fold-change of genes with significant effects

Genotype x site (GxE) interactions were relatively common. In total, 1837 transcripts (corresponding to 19% of expressed genes, Table 1, Table S3) had significant GxE interactions, where the magnitude of expression differences between parental ecotypes depended on the field site in which they were grown (Fig. 2A). Only 94 transcripts had crossing reaction norms (green points, Fig. 2B). More often, expression differences between genotypes were significant at one site and non-significant at the other (1137 transcripts, 11%, blue and red points, Fig. 2).

**Fig. 2.**
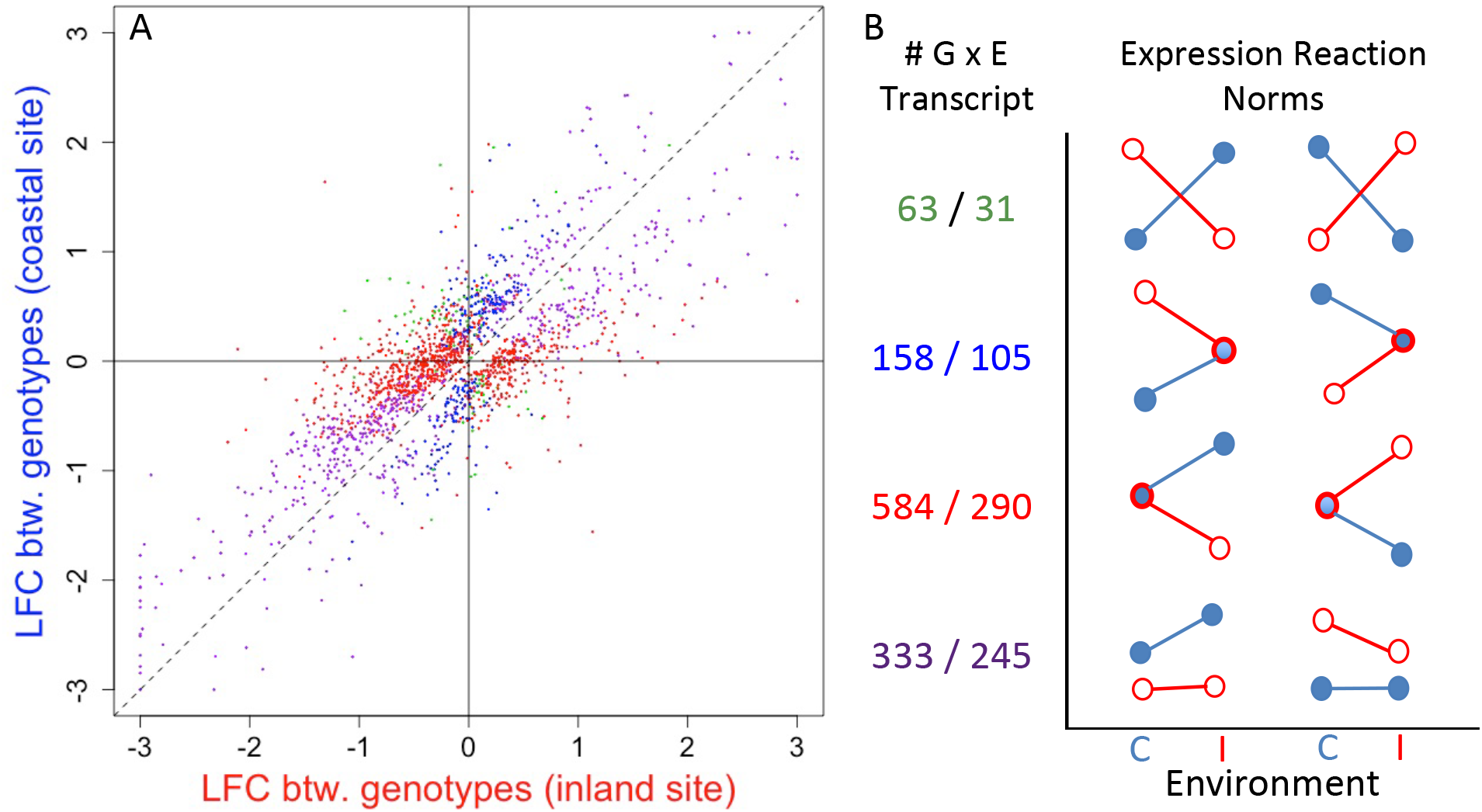
A) Average log2 fold-change (LFC) differences between parental genotypes for expressed transcripts at the two field sites. B) Expression reaction norms for inland (red) and coastal (blue) lines. The numbers correspond to the colored points in part A of the figure. LFC has been restricted to <3 and <−3 for graphing purposes. Note, 28 transcripts with ambiguous expression patterns have been excluded.

Most of the expression differences between the coastal and inland lines could be attributed to *cis*-regulatory variation: 79% (5532 genes) of differentially expressed genes had *cis* regulatory differences, while about half (3611 genes) of the differentially expressed genes had detectable *trans* effects (Fig. 3 & 4, Table 1). At both coastal and inland field sites, the magnitudes of *cis* and *trans* regulatory differences were similar (Table S4). However, we detected far more significant *trans* effects in the inland field site than the coastal field site (Fig. 3 & 4, Table S4). Overall, there was only moderate overlap between the sets of *cis*-regulated transcripts from the two environments (Fig. S7. The same was true for the sets of trans-regulated transcripts from the two environments.

**Fig. 3.**
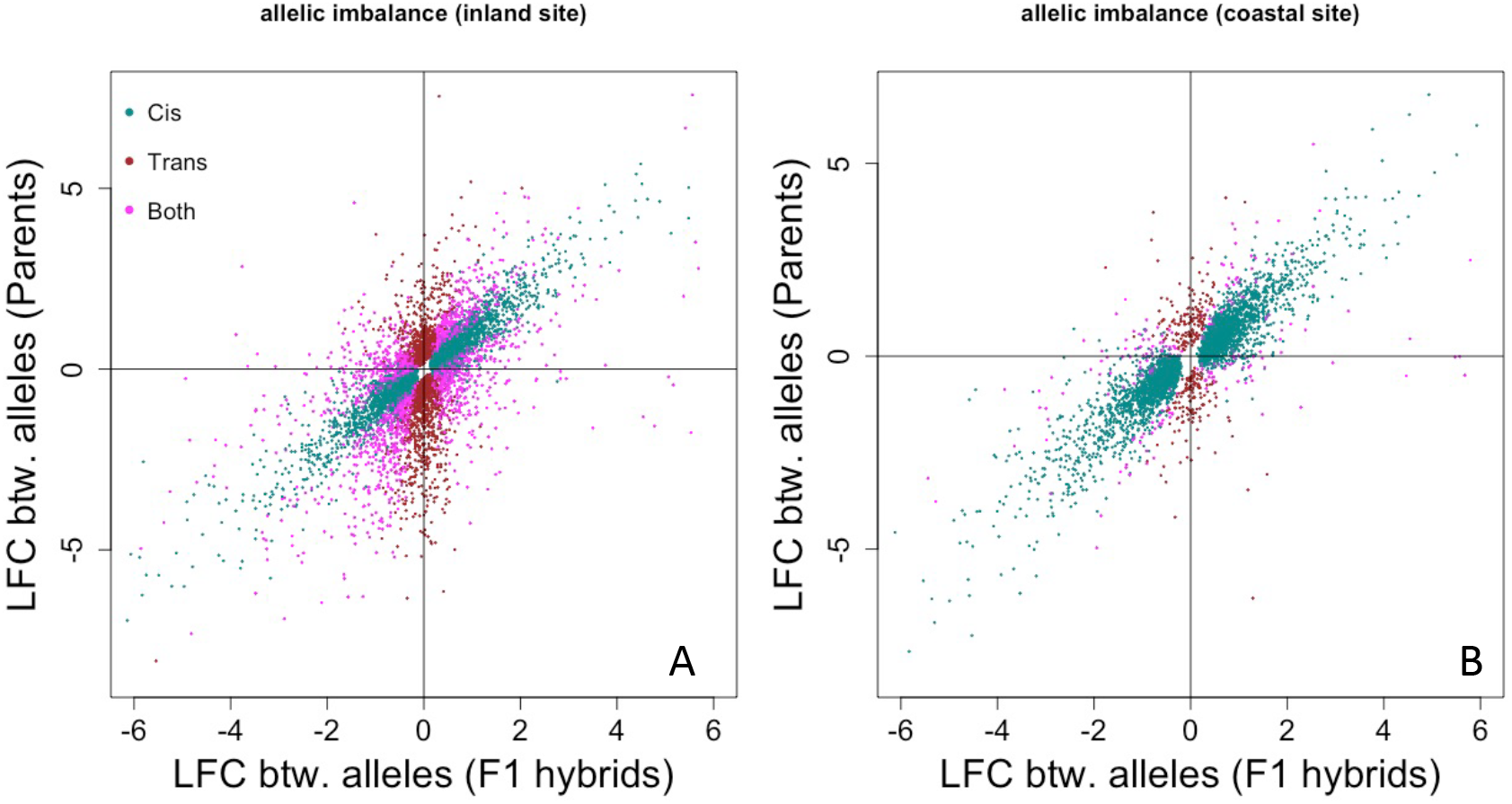
*Cis* and *trans* regulated transcripts detected at the coastal (A) and inland (B) field sites. Average log2-fold change (LFC) ratios are between inland and coastal alleles in the parental (y-axis) and F1 (x-axis) genetic backgrounds.

**Fig. 4.**
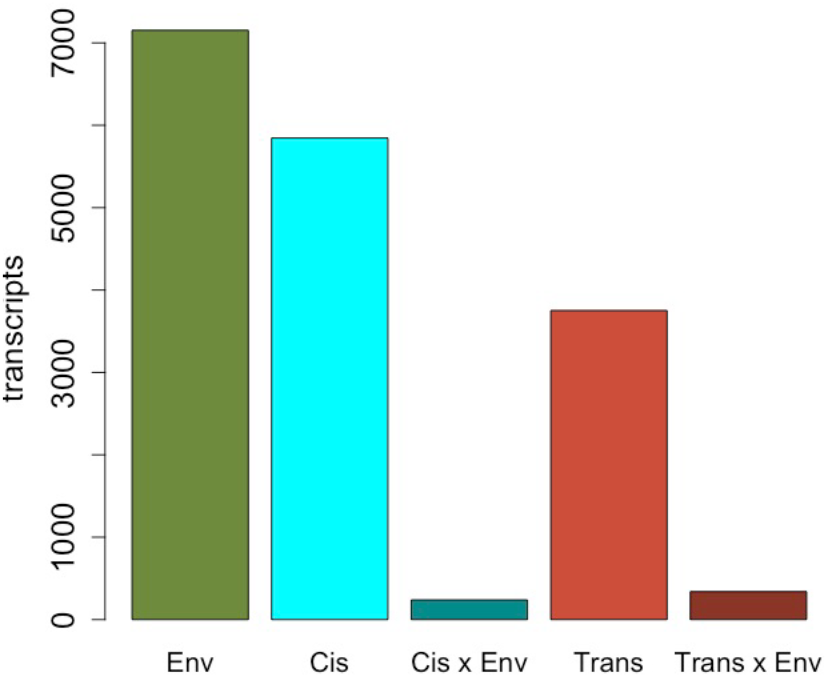
The frequency of significant model effects among all expressed transcripts.

### Gene Ontology Enrichment and Expression Networks

We explored the functional significance of genes with differential expression between environments and between parental ecotypes. Differentially expressed genes in the coastal (SS) parent between sites were enriched for glycosinolate/glucosinolate/S-glycoside biosynthesis and metabolism, response to starvation, and others (Table S1). Differentially expressed genes in the inland parent (LL) between sites were enriched for GO categories related to cuticle development, meristem and floral development, and pigment production. Differentially expressed genes between the parental lines (across sites) were enriched for genes involved with cell wall formation, meristem initiation, and responses to viral pathogens. Genes with genotype x site (GxE) interactions had the most significantly enriched GO terms (*N* = 69), including brassinosteroid biosynthesis/metabolism, cell wall formation, regulation of development, and pigment accumulation.

We used network clustering analysis to examine correlated expression of gene clusters (modules) within habitats for the coastal and inland lines (Pearson’s correlation, *r* > 0.75, Fig. S5. The coastal perennial line (SS) preferentially expressed modules related to root and epidermal development at its home site, while oxidative stress response, chlorophyll metabolism, and circadian rhythm modules were highly expressed in inland habitat (Table S2). The inland annual line preferentially expressed gene clusters related to nucleoside catabolism, sugar and starch metabolism, pigment accumulation, and photoperiodism at the inland field site, but had no modules strongly associated with the coastal field site.

### Chromosomal Inversion Analyses

Chromosomal inversions are thought to act as adaptation supergenes that hold together long haplotypes containing multiple adaptive polymorphisms through suppressed recombination (Kirkpatrick 2010; Joron et al. 2011; Thompson & Jiggins 2014). We were thus interested in evaluating the hypothesis of whether the known inversions in this system hold together haplotypes of differentially expressed genes. There were significantly more differentially expressed genes inside the chromosome 5 inversion than expected by chance and there were significantly more GxE interactions for genes in this inversion (Table 1). The adaptive (Lowry & Willis 2010) chromosome 8 inversion also had significantly more differentially expressed genes, but only at the coastal site (Table 2). Interestingly, the chromosome 8 inversion had a significant enrichment of *cis*-eQTLs (Table 1), suggesting that it may maintain a higher level of regulatory divergence caused by the evolution of *cis*-regulatory elements.

### Patterns of Candidate Gene Expression

To identify genes putatively linked to local adaptation, we cross-referenced our results with a recent genomic outlier analysis, which identified 667 genes with a high level of allelic differentiation between coastal and inland populations (Lowry et al. 2010). We found that the magnitude of differential expression between environments was significantly elevated for candidate genes with highly differentiated SNPs (from Gould et al. 2017) in their promoter regions (*P* < 0.05, Table S5). The magnitude of differential expression between genotypes for candidate genes with differentiated promoters was also higher than for non-candidate genes, but not significantly so (*P* = 0.08). Candidate genes with differentiated gene regions tended to have expression differences between lines or between environments that were lower in magnitude than non-candidate genes.

We were able to examine gene expression in seven top candidate genes highlighted by the outlier analysis and/or previous QTL mapping studies (Fig. 5). The candidate gene *ABA1* codes for the first protein in the biosynthesis of the hormone abscisic acid, which controls responses to osmotic stress, including drought (Xiong et al. 2002). An *ABA1* (abscisic acid deficient 1) ortholog, Migut.H00431, is located within the chromosome 8 inversion and may contribute to inland ecotype’s adaptation to low soil water availability. *ABA1* was always more highly expressed in the inland line and was upregulated in both lines at the inland field site (Fig. 5A). Previously, we showed that coastal *M. guttatus* populations have evolved leaf tissue tolerance to oceanic salt spray (Lowry et al. 2009). Leaf salt tolerance is generally mediated by a Na^+^/H^+^ antiporter (*SOS1*; Qiu et al. 2002), which is potentially activated by a calcium sensor (*CBL10*; Kim et al. 2007). Both *SOS1* (Migut.E00570) and *CBL10* (Migut.A00138) were more highly expressed in the coastal line than the inland line across habitats (Fig. 5B, C), a pattern consistent with elevated salt tolerance of coastal populations.

**Fig. 5.**
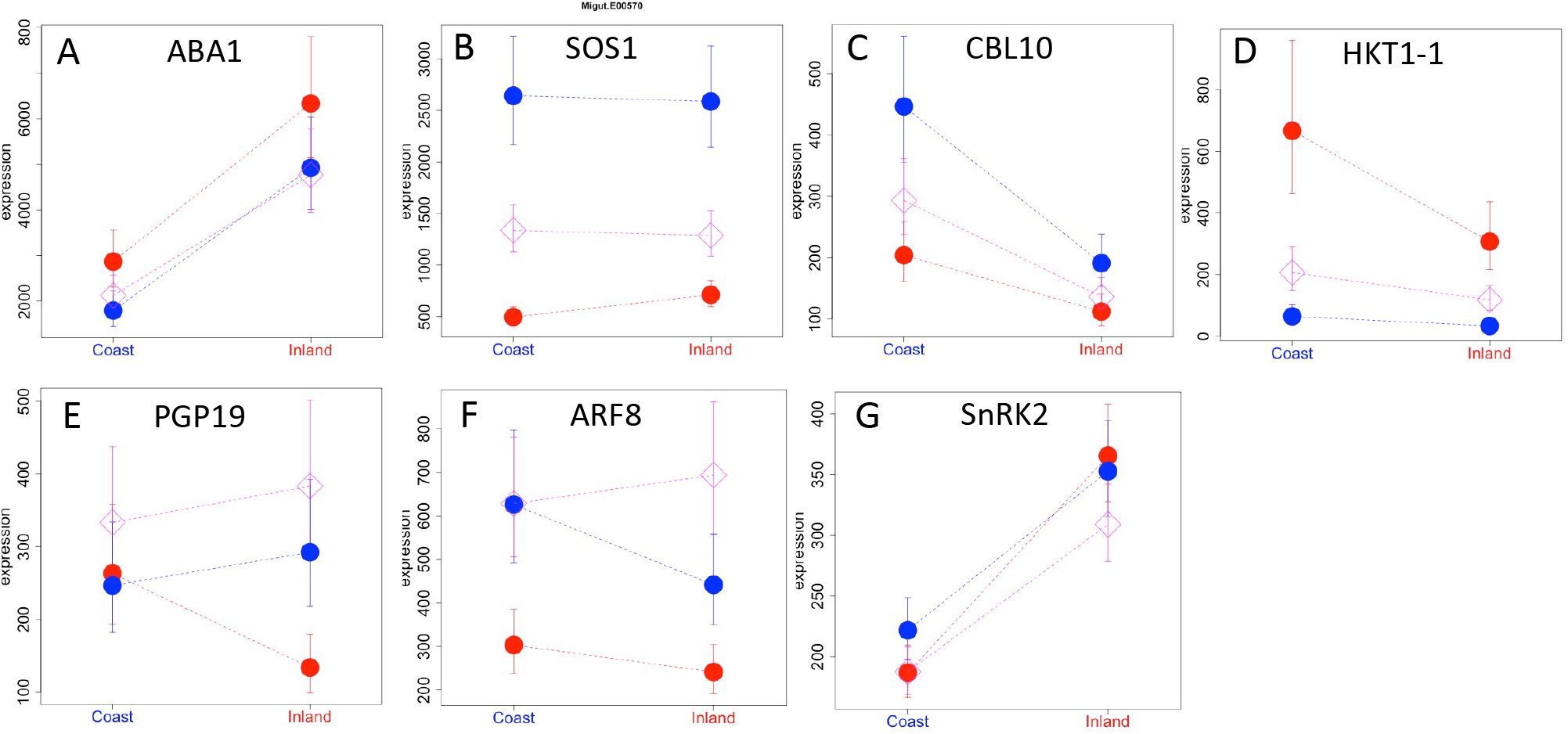
Reaction norms of expression at two field sites for seven candidate adaptive genes. A) ABA1 – abscisic acid deficient 1; B) SOS1 – salt overly-sensitive 1; C) CBL10 -calcineurin B-like protein 10; D) HKT1-1, high-affinity K+ transporter (functions in salt tolerance); E) PGP19, p-glycoprotein 19 (functions in auxin transport); F) ARF8 – auxin response factor 8; G) SnRK2, Snf-related protein kinase (functions in salt tolerance); Red, inland genotype. Blue, coastal genotype. Pink, F1 hybrid.

## DISCUSSION

Overall, our results identified multiple key patterns regarding the relationship between gene expression and the evolution of local adaptation. Differential expression is substantial between locally-adapted accessions in nature. Expression plasticity of alleles across habitats (genotype x environment interactions) appear to be common and may minimize fitness trade-offs at individual loci that contribute to the overall pattern of local adaptation. A large portion of that differential expression is due to *cis*-regulatory divergence. Chromosomal inversions appear to act as supergenes by holding together haplotypes of differentially expressed genes, but this pattern depends on habitat context. We discuss each of these major findings as well as the limitations and caveats associated with our study in detail below.

### Sources of Variation in Gene Expression

One of the major outstanding questions in local adaptation research is why the fitness effects of individual loci on local adaptation are so often asymmetric across habitats. Theoretical models (Levene 1953; Hedrick 1976, 1986; Dieckmann & Doebeli 1999) and studies in key model systems (Abzhanov et al. 2004; Hoekstra et al. 2006; Barrett et al. 2009) have suggested trade-offs at individual loci (e.g. antagonistic pleiotropy) to be common. However, field reciprocal transplant experiments and experimental evolution studies have often found a pattern of conditional neutrality, where loci have a strong effect on fitness in one environment but have an undetectable effect on fitness in alternative environments (reviewed in Bono et al. 2017; Wadgymar et al. 2017). This pattern has also been found in the coastal perennial/inland annual *M. guttatus* system, where three salt tolerance QTLs have fitness effects in coastal habitat but no detectable effects in inland habitat (Lowry et al. 2009). While a failure to statistically detect a trade-off across habitats does not rule out the possibility of a small trade-off for an individual locus, it does suggest that adaptive loci are often strongly asymmetric in their fitness effects across habitats.

Gene regulatory divergence between locally adapted populations is one possible mechanism by which trade-offs could be minimized across habitats for individual loci (Rutherford 2003; Ghalambor et al. 2007; Des Marais et al. 2013, 2017; Dayan et al. 2015; Lohman et al. 2017). While some gene regulatory differences between ecotypes may evolve due to genetic drift in isolated populations, adaptive phenotypic plasticity can evolve due to natural selection. Adaptations of organisms to maintain homeostasis when faced by environmental challenges can be caused by genotype specific changes in gene expression (Gibson 2008; Des Marais et al. 2017). However, such environmentally dependent changes in gene expression would not be expected to result in a cost if gene expression did not differ among genotypes in habitats that lacked particular environmental stresses. In our study, we identified 1137 transcripts that were differentially expressed between the coastal perennial and inland annual lines in one habitat (coast or inland) but did not show the same expression difference in the alternative habitat (GxE transcripts, Fig. 2). In contrast, fewer transcripts (733) had the same expression difference between ecotypes regardless of environment (G-only transcripts). For the current experiment, we cannot definitively determine whether any of the GxE or G-only transcripts cause asymmetries in fitness effects across habitats. Further, some of the genotype specific changes in gene expression could potentially be maladaptive (Ghalambor et al. 2007). Even so, our results do clearly show that habitat-specific differential gene expression is common for locally adapted ecotypes. Future studies should focus on making the critical linkages between patterns of gene expression and fitness in the field to determine whether asymmetries of fitness effects across habitats are the result of habitat-specific differential gene expression.

Overall, *cis*-regulatory evolution was responsible for at least 79% of the differentially expressed genes. This is about twice the percentage of *cis*-regulated differentially expressed genes for crosses both within and between *Drosophilia* species (Coolon et al. 2014). However, the relatively high proportion of *cis*-regulatory divergence underlying differentially expressed genes is not too surprising, given the high level of polymorphism among *M. guttatus* accessions (Brandvain et al. 2014; Gould et al. 2017; Puzey et al. 2017). Further, in terms of the proportion of *cis* regulated genes out of all genes surveyed (59% for our study), our results are not very different from other study systems. Mack et al. (2016) found significant *cis* effects for 68% of surveyed genes in crosses between house mice subspecies (*Mus musculus musculus* and *M. m. domesticus*). Steige et al. (2015) found that 33-39% of genes surveyed had significant allele-specific expression effects for hybrids between accessions of the plant *Capsella grandiflora*.

One striking pattern we observed in this study was a great deal more differential regulation between ecotypes in the inland environment than the coastal environment (Table 2, Fig.3). We can attribute only a small part of this difference to greater experimental power to detect differences at the inland site (75% inland vs. 68% coast, for transcripts with a minimum fold-change of 2X). Because mapping bias is well controlled in this study, we do not believe this overall pattern is the result of an analytical artifact. We hypothesize that part of this difference is due to more stressful conditions experienced by plants at the inland site; plants that survive to maturity in coastal habitats are generally larger than plants growing at the inland habitat (Lowry et al. 2008; Popovic & Lowry *unpublished data*). It remains to be seen whether the magnitude of gene regulatory divergence between ecotypes varies across habitats in other systems as well.

### Gene Regulatory Evolution and Chromosomal Inversions

Chromosomal inversions have been thought to be involved in evolutionary adaptations since classic studies in *Drosophila* found that inversion polymorphisms are frequently correlated with environmental conditions over geographic space and through time (Dobzhansky 1951, 1970; Kirpatrick & Barton 2006; Hoffman & Rieseberg 2008; Cheng et al. 2012; Ayala et al. 2013, 2014; Adrion et al. 2015). Inversions strongly suppress genetic recombination in heterokaryotic (inversion heterozygotes) individuals because recombinant gametes are unbalanced (Dobzhansky 1970; Rieseberg 2001). Researchers have thus hypothesized that inversions could act as supergenes by holding together long haplotypes containing multiple adaptive polymorphisms through suppressed recombination (Dobzhansky 1970; Joron et al. 2011; Thompson & Jiggins 2014; Kirkpatrick & Barton 2006; Kirkpatrick & Barrett 2015).

The supergene hypothesis is a major reason why inversions are thought to frequently contribute to the evolution of adaptations (Dobzhansky 1970; Kirkpatrick & Barton 2006; Hoffman & Rieseberg 2008; Faria & Navarro 2009; Thompson & Jiggins 2014). Early researchers, especially Dobzhansky (1970), argued that inversions could hold together co-adapted epistatic complexes of genes through suppressed recombination. However, theoreticians have recently shown that inversions can spread due to purely additive effects, when at least two locally adaptive loci are captured within the inverted region (Kirkpatrick & Barton 2006; Kirkpatrick & Barrett 2015). Regardless of the ultimate genetic mechanisms, these models all predict that suppression of recombination by inversions maintains adaptive divergence at multiple linked loci.

Global gene expression analyses offer one way to test whether inversions hold together haplotypes of genetic variation potentially responsible for phenotypic divergence between locally adapted populations. Research in a few systems has found evidence of increased levels of expression divergence in inversions (Marquès-Bonet et al. 2004; Cassone et al. 2011; Fuller et al. 2016). For example, Marquès-Bonet et al. (2004) found 8.9% greater divergence (fold change) in human versus chimp brain gene expression for chromosomal inversions when compared to the rest of the genome. We did not observe a significant difference in fold change between inverted and non-inverted regions in our study. However, genes within the region of the chromosome 5 inversion were 12.5% more likely to be differentially expressed. Genes within the adaptive chromosome 8 inversion were 8.3% more likely to have a *cis*-eQTL.

The enrichment of gene expression differences in our study is consistent with patterns found in inversions between chimpanzees and humans (Marquès-Bonet et al. 2004) and *Drosophila* (Fuller et al. 2016; Said et al. 2018), but contrasts with results for the plant *Boechera stricta* (Lee et al. 2017). While the finding of an enrichment of differential gene expression for inversions in this study and others is consistent with the supergene hypothesis of inversion evolution, it is by no means definitive proof. Demonstrating that an inversion has evolved as an adaptation supergene requires confirmation that multiple genes within the inversion have effects on adaptive phenotypes (Hoffman & Rieseberg 2008; Kirkpatrick 2010; Kirkpatrick & Kern 2012). There is currently no study that we are aware of which has experimentally confirmed that two or more genes within an inversion contribute additively or epistatically to adaptive phenotypic differences between locally adapted populations, ecotypes, or species (but see Kunte et al. 2014).

### Candidate Genes and Pathways

Analysis of expression across both ecotypes and environments allowed us to examine the detailed expression patterns of several candidate genes that may contribute to adaptive divergence. We observed a diversity of expression patterns across candidate genes (Fig. 5). The gene *ABA1* catalyzes the first step in the synthesis of the abiotic stress hormone ABA, and the *Mimulus* ortholog of this gene is located in the adaptive chromosomal inversion on chromosome 8. In *Arabidopsis* and other plants, increased ABA production leads to drought tolerance by triggering stomatal closure and resistance to osmotic stress (Jakab et al 2005). Previous work has shown that the chromosome 8 inversion in *M. guttatus* contributes directly to fitness differences between coastal and inland lines (Lowry & Willis 2010) and that the genic region of *ABA1* within the inversion has an unusually high degree of differentiation (high *FST*) between coastal and inland populations (Gould et al. 2017). In the present study, we found that both genotypic and environmental differential regulation contribute to expression patterns at this locus. *ABA1* was always more highly expressed in the inland ecotype and also upregulated in the inland environment by both ecotypes (Fig. 5A). In contrast, the candidate gene P-glycoprotein19, *PGP19* (Migut.H00909), also located in the adaptive chromosome 8 inversion (Gould et al. 2017), was differentially regulated between ecotypes only at the inland field site (Fig. 5E). The *Arabidopsis* ortholog of *PGP19* is an ATP transporter involved in gravitropism, growth, and development by mediating auxin signaling (Okamoto et al 2015).

Consistent with oceanic salt spray playing a key role in local adaptation (Lowry et al. 2009), multiple salt tolerance candidate genes were differentially regulated between the coastal perennial and inland annual ecotypes. Two major genes in the Salt Overly Sensitive (*SOS*) pathway, *CBL10* and *SOS1*, were expressed at higher levels in the coastal ecotype than the inland ecotype across habitats (Fig. 5B, C). *CBL10* is a calcineurin B-like calcium sensor that interacts with the protein kinase *SOS2* in the shoots of plants to positively regulate Na^+^ transporters in the tonoplast of the vacuole as well as potentially in the plasma membrane (Quan et al. 2007; Kang & Nam 2016; Egea et al. 2018). The function of *CBL10* in salt response has been shown to be conserved across *Arabidopsis* (Quan et al. 2007), *Populus* (Tang et al. 2014), and tomatoes (Egea et al. 2018). Interestingly, *CBL10* has been found to confer salt tolerance in plants, while at the same time making them more susceptible to drought (Kang & Nam 2016), suggesting a potential trade-off for local adaptation to coastal habitats that are inundated by salt spray. *SOS1* is a well characterized Na^+^/K^+^ antiporter that pumps toxic Na^+^ ions out of the cytoplasm of plant cells into the surrounding apoplast (Shi et al. 2002). *SOS1* is potentially positively regulated by the *CBL10/SOS2* complex in the shoots of plants (Quan et al. 2007). In addition to the *SOS* pathway, there were G and E effects (but not GxE) for an ortholog of the High-Affinity K+ Transporter1, *HKT1. HKT1* has previously been linked in the evolution of salt tolerance adaptation in coastal populations of *Arabidopsis thaliana* (Fig. 5D; Rus et al. 2006; Baxter et al. 2010).

### Caveats, Limitations, and Future Directions

There are important caveats and concerns to consider in the interpretation of the results of our study. One of the foremost caveats of the study is its narrow scope. We only evaluated gene expression differences between one inbred line per ecotype, at two field sites, and for gene expression at one time point. Despite these drawbacks, this is the first study that we are aware of to evaluate allele-specific expression in the field between known locally adapted populations (Voelckel et al. 2017), which allowed us to detect novel *cis* x habitat and *trans* x habitat interaction (Fig. 4; Table S6). The major outstanding questions include how variable gene expression is across populations, field sites, and tissue types? In the future, it will be critical to evaluate gene expression at different time points and across tissue types to gain a more comprehensive understanding of how ecotypic differences in gene expression manifest throughout development under natural field conditions.

Perhaps, the greatest challenge for interpreting gene expression results from reciprocal transplant experiments is understanding how patterns of expression actually contribute to phenotypes and fitness across habitats. Fitness trade-offs that are associated with local adaptation could potentially be caused by any type of gene expression pattern. Constitutive expression differences between ecotypes could cause stable phenotypic differences that are beneficial in one habitat but deleterious in another. Gene expression characterized by genotype x environment interactions could cause fitness trade-offs, but just as likely could mitigate trade-offs by helping ecotypes maintain homeostasis across habitats.

Until recently, studies of gene regulation have for the most part been limited to the laboratory setting. Coupling those studies with reciprocal transplant experiments to better understand local adaptation is still rare (Kenkel & Matz 2016; Lohman et al. 2017; Voelckel et al. 2017). Allele-specific expression is clearly an important tool for understanding the evolution of regulatory differences between locally adapted populations, ecotypes, and species. However, we hope that future reciprocal transplant experiments will go beyond RNA-sequencing by also examining genotypic differences in genome-wide patterns of chromatin remodeling and protein translation across divergent habitats (Rodgers-Melnick et al. 2016; Voelckel et al. 2017).

## DATA ACCSSIBILITY

The assembled reference transcriptome and normalized count data for all transcripts in all individuals has been archived at Data Dryad (see DOI: https://doi.org/10.5061/dryad.57d6118).

## ACKNOWLEDGMENTS

We thank D. Des Marais, J. Tung, J. Puzey, J. Friedman, R. Guerrero, R. Rellán-Álvarez, J. Lasky, and six anonymous reviewers for comments that helped to improve our manuscript. We thank K. Childs and J. Lovell for guidance with expression analyses, B. Blackman and E. Patterson for starting seeds for us at UC Berkeley, and D. Popovic for assistance in field planting. The Bodega Marine Reserve and the Pepperwood Preserve kindly allowed us to conduct this reciprocal transplant field experiment. Funding was provided by Michigan State University through a start-up package to DBL. The authors declare no conflicts of interest.

## DATA ACCESSIBILITY

RNA-sequence data are deposited are available through the NCBI Sequence Read Archive under project identifier XX.

## AUTHOR CONTRIBUTIONS

DBL designed and conducted the field experiment. YC conducted the laboratory work. BAG analyzed the data. DBL and BAG wrote the manuscript.

